# Oligodendrocyte Enriched Brain Organoids Reveal Impaired Oligodendroglial Maturation and Altered Neural Network Activity in Down Syndrome

**DOI:** 10.64898/2026.08.27.747500

**Authors:** Marta Boira Marti, Sean D. Morrison, Selin Pars, Bahaa Al-mhanawi, Peter G. Noakes, Ernst J. Wolvetang, Mohammed R. Shaker

## Abstract

Individuals with Down syndrome (DS) display developmental delay, intellectual disability, premature brain ageing, and an increased risk of Alzheimer-like neurodegeneration. Although the neuropathology of the postnatal and adult DS brain has been widely described, it remains unclear how trisomy 21 alters early human neural and glial development. Here, we used human oligodendrocyte enriched brain organoids derived from trisomic and euploid iPSCs to define the cellular, functional, and molecular consequences of trisomy 21 during early brain development. Trisomic organoids exhibited an early growth delay, reduced oligodendroglial specification, and impaired oligodendrocyte maturation, resulting in decreased myelination. These defects were accompanied by increased astroglial output and delayed neuronal maturation. At the functional level, trisomic organoids showed elevated spontaneous network activity, but failed to mount normal coordinated responses to pharmacological stimulation, consistent with abnormal neural circuit development. Bulk RNA sequencing revealed the strongest transcriptomic dysregulation occurs at early neural and glial specification. Together, these findings show that trisomy 21 disrupts early developmental stages and establish oligodendrocyte enriched brain organoids as a human model to investigate the developmental origins of white matter and network dysfunction in DS.

## Introduction

Down syndrome (DS) is the most frequent viable chromosomal aneuploidy, arising from trisomy of human chromosome 21 (HSA21) and affecting ∼1 in 700 live births [1]. In addition to systemic manifestations, trisomy 21 consistently disrupts central nervous system (CNS) development. Neuropathological hallmarks include microcephaly, hypocellularity of neurons and oligodendrocytes (OLs), defective dendritic arborisation, perturbed synaptic morphology, early-onset Alzheimer-like pathology and premature brain ageing [2,3]. Diffusion-tensor imaging and post-mortem analyses further reveal widespread white-matter abnormalities and persistent hypomyelination that correlate with cognitive and motor dysfunction [4,5].

Mouse models carrying partial or complete triplication of HSA21-orthologous regions, most notably Ts65Dn, have been instrumental in dissecting DS neuropathology. These studies demonstrate intrinsic defects in OL differentiation and myelin production that slow axonal conduction, accompanied by transcriptomic down-regulation of key myelination genes (e.g., *CNP*, *PLP1*, *SOX10*, *GPR17*, *C21ORF91*) during hippocampal development [6]. In parallel, Ts65Dn mice exhibit excessive GABAergic interneuron output, altered excitatory/inhibitory balance, abnormal astrocyte overproduction and enhanced microglial activation, highlighting the contribution of non-cell-autonomous mechanisms to white-matter pathology [7]. While invaluable, species-specific differences in gene dosage, developmental timing and myelin composition limit the translatability of these findings to the human condition.

Human induced pluripotent stem cells (iPSCs) have provided a complementary, patient-specific platform to model DS. Early two-dimensional iPSC studies reproduced decreased neurogenesis, reduced NPC proliferation and altered DSCAM-PAK1 signalling, but could not capture higher-order cytoarchitecture [8,9]. Three-dimensional (3D) cerebral organoids overcome this limitation by self-organising into region-specific domains that recapitulate cell-cell and cell-matrix interactions [10]. Our laboratory recently established choroid-plexus-cortical organoids that faithfully mirrored DS-associated epithelial and neuronal defects and revealed enhanced neurotropism of SARS-CoV-2 [11]. Nevertheless, conventional brain organoids generate mature, myelinating OLs only after prolonged (>110 day) culture, hindering systematic interrogation of hypomyelination.

To address this knowledge gap, our lab has developed an oligodendrocyte-enriched brain organoid (OLBO) protocol that yields myelinating OLs alongside physiologically maturing neurons, astrocytes and microglia within 42 days [12–14]. Here we deploy DS and control OLBOs to dissect both cell-autonomous OL deficits and the influence of neighbouring neural and glial populations on myelination. Using quantitative immunohistochemistry and bulk RNA-sequencing, we identify molecular and cellular drivers of hypomyelination in trisomy 21. Our findings underscore the utility of accelerated OLBOs as a tractable human model for delineating DS white-matter pathology and for future therapeutic screening.

## Results

### Generation and Characterisation of Oligodendrocyte-Enriched Brain Organoids

We recently established a robust protocol for generating oligodendrocyte-enriched brain organoids (OLBOs) that recapitulate key cellular features of human CNS development, including functional oligodendrocytes, neurons, and astrocytes [12,14]. Human iPSCs were initially directed toward a neuroectodermal fate using dual SMAD inhibition, resulting in the formation of two-dimensional human neuroectodermal (hNEct) colonies within three days. These colonies were subsequently transitioned into three-dimensional neural spheroids and expanded under FGF2 stimulation to promote neural progenitor proliferation (Fig. 1A). To accelerate neuroepithelial polarisation and oligodendroglial lineage specification, spheroids were embedded in low concentrations of Matrigel and exposed from day 3 onward to a defined combination of oligodendrocyte-promoting factors, including PDGF-AA, HGF, IGF-1, T3, and NT-3. Concurrent supplementation with cAMP, B27 without vitamin A, biotin, and β-mercaptoethanol supported oligodendrocyte progenitor cell (OPC) survival and maturation, while N2, insulin, and non-essential amino acids enabled co-differentiation of neurons and astrocytes (Fig. 1A). This stepwise strategy yielded OLBOs within a markedly shortened timeframe compared with conventional cerebral organoid protocols.

**Figure 1.**
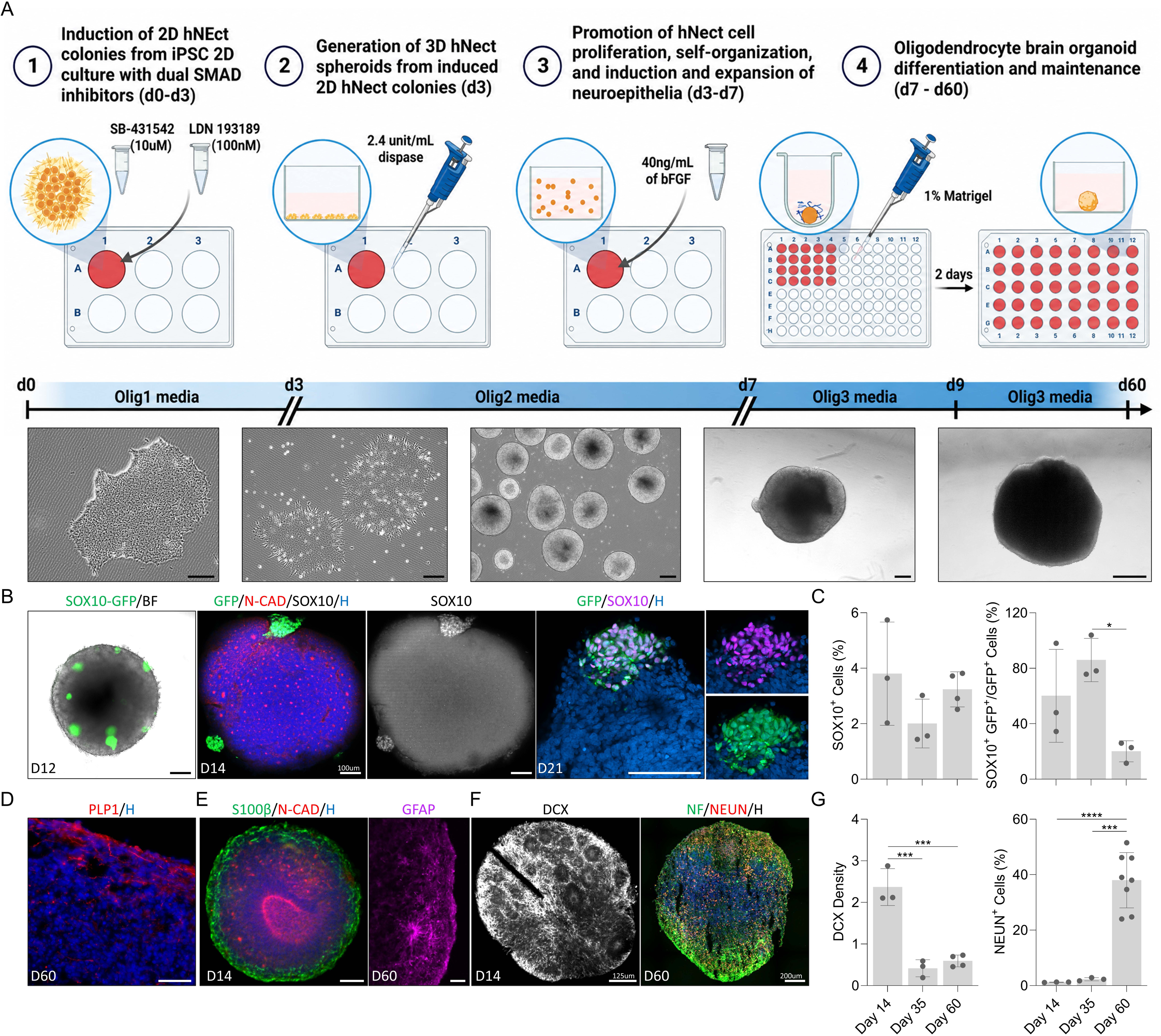
Generation and multilineage validation of oligodendrocyte enriched brain organoids. **(A)** Schematic of the four-step protocol used to direct human iPSCs into 3D oligodendrocyte-enriched brain organoids. Below are representative brightfield/phase-contrast images show iPSCs and organoids morphology over the course of differentiation. Scale bars, 200 µm. **(B)** Brightfield image of neural spheroid at day 12 derived from SOX10-GFP reporter hiPSC (scale bar, 200 µm), and wholemount immunostaining images of neural spheroids stained with oligodendroglial lineage (SOX10) as well as neural rosettes marker (N-cadherin), scale bar = 100 µm. Nuclei counterstained with Hoechst (Blue). **(C)** Bar graphs show the quantification of oligodendroglial cells in OL-brain organoids over time with percentage of total cells expressing SOX10 (left panel) and the proportion of SOX10⁺/GFP⁺ co-labelled cells relative to the total GFP⁺ lineage (right panel) at days 14, 35, and 60. Data are means ± S.D.; n = 19. \**P* < 0.05, via one-way analysis of variance (ANOVA). **(D)** Confocal image of sectioned OL-brain organoids staining mature oligodendrocytes with PLP1 (red) at day 60. Nuclei counterstained with Hoechst (blue). Scale bar = 50 µm. **(E)** Wholemount immunostaining of astroglial specification and maturation marked with the early glial marker S100β (green) and neural rosettes N-CAD (red) at day 14, scale bar = 50 µm. Left panel is immunostaining image of sectioned OL-brain organoids stained with GFAP (magenta) at day 60, nuclei counterstained with Hoechst (H), scale bar = 50 µm. **(F)** Wholemount immunostaining of neuronal lineage progression stained with DCX (Gray) at day 14 (scale bar = 125 µm), and neurofilament (Green) with NEUN (red) at day 60 (scale bar = 200 µm). Nuclei counterstained with Hoechst (Blue). **(G)** Bar graphs show the quantification of neuronal cells in OL-brain organoids over time with density of DCX signal normalised to total area (left panel) and the percentage of NEUN⁺ cells normalised to total cells labelled with Hoechst (right panel) at days 14, 35, and 60. Data are means ± S.D.; n = 24. \*\**P* < 0.01, \*\*\**P* < 0.001, \*\*\*\**P* < 0.0001 via one-way analysis of variance (ANOVA).

To track oligodendroglial lineage emergence and dynamics, we employed a SOX10-GFP reporter hiPSC line [15], in which GFP expression is restricted to oligodendroglial cells. Discrete clusters of GFP-positive cells became detectable at the organoid periphery as early as day 12 (Fig. 1B), consistent with the formation of OPC-enriched niches. Immunohistochemical analysis confirmed that these GFP-positive clusters corresponded to SOX10-expressing cells, a transcription factor essential for oligodendroglial specification and differentiation. SOX10-positive cells were already evident by day 7, accumulated at the organoid edges during early development, and progressively dispersed across the organoid surface from day 35 onward, coinciding with oligodendrocyte maturation (Fig. S1A). Quantitative analysis revealed dynamic changes in oligodendroglial populations over time, approximately 4% of cells were SOX10-positive at day 14, decreasing to ∼2% by day 35 and stabilising at ∼3% by day 60 (Fig. 1C). Co-expression analysis showed that SOX10 and GFP overlapped in over 60% of cells at day 14 and more than 80% at day 35, reflecting active OPC proliferation during early stages. By contrast, only ∼20% of cells co-expressed SOX10 and GFP at day 60 (Fig. 1C, S1B), indicating lineage diversification and maturation. These findings suggest that early SOX10-positive OPCs give rise to both mature oligodendrocytes and alternative glial lineages over time. Consistent with oligodendrocyte maturation, OLBOs at day 60 expressed late-stage oligodendrocyte markers, including PLP1 and CNPase, indicative of the transition from progenitor to differentiated states (Figs. 1D, S1C). Fully differentiated, myelinating oligodendrocytes were further confirmed by robust expression of myelin basic protein (MBP), localised to membranous structures within the organoids (Fig. S1C). Together, these data demonstrate efficient and accelerated generation of mature oligodendrocytes within OLBOs.

Astrocyte specification was assessed using S100β and GFAP as early and mature astrocytic markers, respectively. S100β-positive cells emerged as early as day 7 and were detected both within peripheral OPC niches and central regions by day 14 (Fig. 1E). From day 35 onward, S100β-positive cells dispersed throughout the organoid. Notably, S100β expression only partially overlapped with GFP, and not all GFP-positive cells are S100β -positive cells (Fig. S1D), consistent with previous reports indicating that S100β is not exclusive to astrocytes and can also mark OPCs [16]. GFAP-positive mature astrocytes became apparent from approximately day 60, confirming progressive astrocytic maturation within OLBOs.

To evaluate neuronal development and maturation, we examined the expression of doublecortin (DCX), a marker of immature neurons. DCX-positive cells were detected by day 7 and exhibited a widespread distribution by day 14 (Fig. 1F). As development progressed, DCX-positive progenitors reorganised and became enriched at peripheral regions, forming defined neural progenitor pools (Fig. S1E). Quantification revealed a significant, time-dependent reduction in DCX density, indicating neuronal maturation (Fig. 1G). In parallel, expression of the mature neuronal marker NEUN was detectable from day 14 and increased significantly by day 60, rising from approximately 2% to nearly 7% of total cells (Figs. 1G, S1F).

Collectively, these findings demonstrate that the OLBO protocol supports rapid and coordinated differentiation of oligodendrocytes alongside neurons and astrocytes. The early establishment of OPC niches, followed by oligodendrocyte maturation and integration with neuronal networks, provides a physiologically relevant and temporally accelerated platform for investigating human myelination and white matter pathology.

### Trisomic OL-Brain Organoids Recapitulate Key Neurodevelopmental Abnormalities Associated with Down Syndrome

To determine whether OLBOs recapitulate key features of DS neuropathology, we generated organoids from a trisomic iPSC line (DS18) and a euploid control line (EU79) and compared their growth and lineage progression over time (Fig. 2A). Consistent with previous reports of reduced brain size and impaired neurodevelopment in DS, trisomic DS18 OLBOs displayed a clear growth deficit during early development. Although both groups increased in size over time, DS18 organoids were already smaller at early stages and remained significantly reduced in diameter from approximately day 14 to day 35 relative to EU79 organoids (Fig. 2A). This difference gradually diminished thereafter, and by day 42 the size of trisomic DS18 organoids approached that of euploid controls (Fig. 2A). These findings indicate that trisomy 21 primarily affects early organoid expansion and patterning rather than later overall tissue growth.

**Figure 2.**
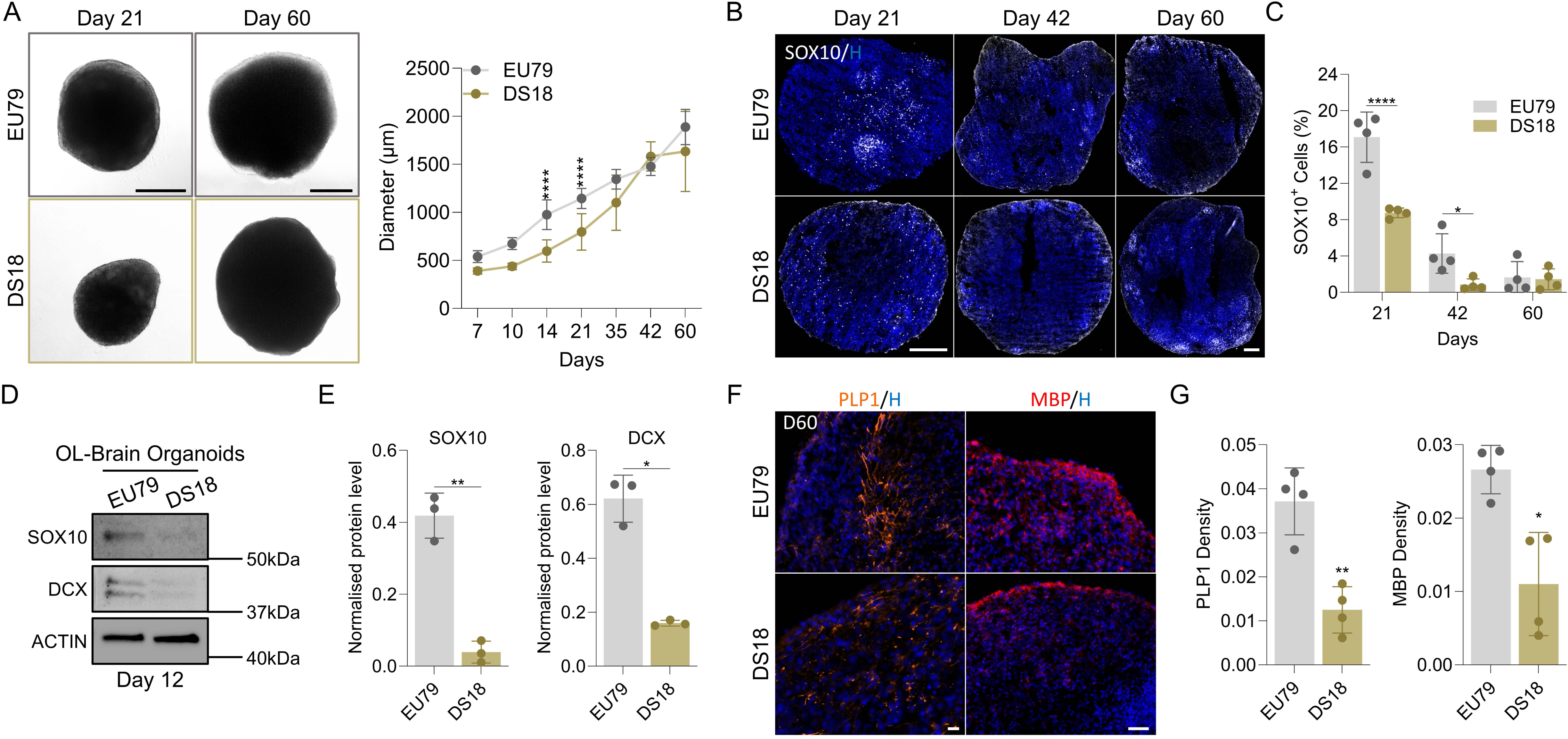
Trisomy 21 impairs oligodendroglial specification and myelin marker expression in OL brain organoids. **(A)** Images of brightfield morphology and growth kinetics of euploid (EU79) and trisomic (DS18) OL-brain organoids at day 21 and day 60 (scale bars = 500 µm). Right panel graph is showing growth of OL-brain organoids in both DS18 and EU79 groups over days 7-60. **(B)** Images of sectioned OL-brain organoids stained with SOX10 (Gray) and nuclei counterstained with Hoechst (Blue) in EU79 and DS18 groups at days 21, 42, and 60. Scale bar = 200 µm. **(C)** Bar graph show the quantification of percentage of SOX10 positive cells in OL-brain organoids over time normalised to total nuclei at days 21, 42, and 60. Data are means ± S.D.; n = 24. \**P* < 0.05, \*\*\*\**P* < 0.0001 via unpaired two tailed Student’s *t* test. **(D)** Western blots showing the levels of SOX10, DCX, and ACTIN in day 12 OL-brain organoids. **(E)** Bar graphs show the quantification of SOX10, as well as DCX levels obtained from **(D)**. Data are shown as means ± S.D. The number of independent experiments = 3, total number of examined organoids is 36. **(F)** Images of sectioned OL-brain organoids stained with PLP1 (Orange), MBP (Red) and nuclei counterstained with Hoechst (Blue) in EU79 and DS18 groups at day 60. Scale bar = 50 µm. (**G**) Bar graphs show the quantification of density of PLP1 signal intensity and MBP signal intensity normalised to total area at day 60 in OL-brain organoids. Data are means ± S.D.; n = 16 \**P* < 0.05, \*\**P* < 0.01 via unpaired two tailed Student’s *t* test.

Because white matter abnormalities and oligodendrocyte deficits are prominent features of the DS brain, we next examined oligodendroglial development in trisomic OLBOs. Immunostaining across developmental stages, which showed a lower abundance of SOX10-positive cells in trisomic DS18 organoids during the early and intermediate phases of differentiation (Fig. 2B). Quantification demonstrated that SOX10-positive cells were significantly reduced in DS18 organoids at day 21 and day 42 compared with EU79 organoids, whereas by day 60 the difference was no longer apparent (Fig. 2C). This finding was confirmed by immunoblot analysis at day 12, which revealed reduced SOX10 protein levels in DS18 organoids relative to euploid controls (Figs. 2D-E), suggesting impaired early oligodendroglial specification. Thus, trisomic OLBOs exhibit a delay or deficit in early oligodendroglial lineage establishment.

We next assessed whether this early impairment was associated with defective oligodendrocyte maturation. At day 60, both trisomic and euploid organoids expressed the late oligodendrocyte marker PLP1, indicating that oligodendroglial differentiation does occur in both groups (Fig. 2F). However, PLP1 signal intensity was reduced in trisomic DS18 organoids, the mature myelinating oligodendrocyte marker MBP was also markedly decreased in DS18 OLBOs relative to controls (Fig. 2F). Quantitative analysis confirmed a significant reduction in both PLP1 and MBP in trisomic DS18 organoids (Fig. 2G), indicating that although oligodendroglial cells are ultimately generated, their maturation and myelin-producing capacity are compromised. Together, these findings are consistent with hypomyelination arising from impaired oligodendrocyte lineage progression in DS.

Given that OPCs can adopt alternative glial fates [17], and that increased astrogliogenesis has been reported in DS [18–20], we next examined astrocyte development. GFAP-positive cells were first evident around day 42 in both groups but were substantially more abundant in trisomic organoids by day 60 (Fig. 3A). Quantification confirmed a significant increase in GFAP-positive cells in DS18 OLBOs at the later stage examined (Fig. 3B). These data support a shift in glial lineage output in trisomic DS18 organoids, favouring astroglial differentiation at the expense of oligodendroglial maturation. We then asked whether neuronal development was also altered in trisomic DS18 OLBOs. Immunostaining for DCX, a marker of immature neurons and early neuronal progenitors, showed its presence in both groups at early stages of differentiation (Fig. S2A). However, DCX density was significantly lower in DS18 organoids than in EU79 controls at day 21 (Fig. S2B), suggesting reduced or delayed early neurogenesis. Consistent with these obervations, immunoblotting at day 12 detected lower DCX protein levels in trisomic organoids than in euploid controls (Figs. 2D-E). Taken together, these findings indicate that neuronal lineage progression is already perturbed during the early stages of trisomic OLBO development. To assess later neuronal maturation, we analysed expression of the post-mitotic neuronal marker NEUN. NEUN-positive cells were detected in both groups from day 14 onward (Fig. S2C), but their abundance increased more strongly in euploid organoids over time (Fig. S2C; Fig. 3C-3D). By day 42, EU79 organoids showed a marked enrichment of NEUN-positive cells, whereas trisomic organoids exhibited a significantly weaker increase (Fig. S2C; Fig. 3D). Although NEUN-positive cells further accumulated by day 60 in both groups (Fig. 3C), the neuronal deficit in DS18 organoids persisted relative to controls (Figs. 3C-D). These data indicate delayed and attenuated neuronal maturation in trisomic OLBOs.

**Figure 3.**
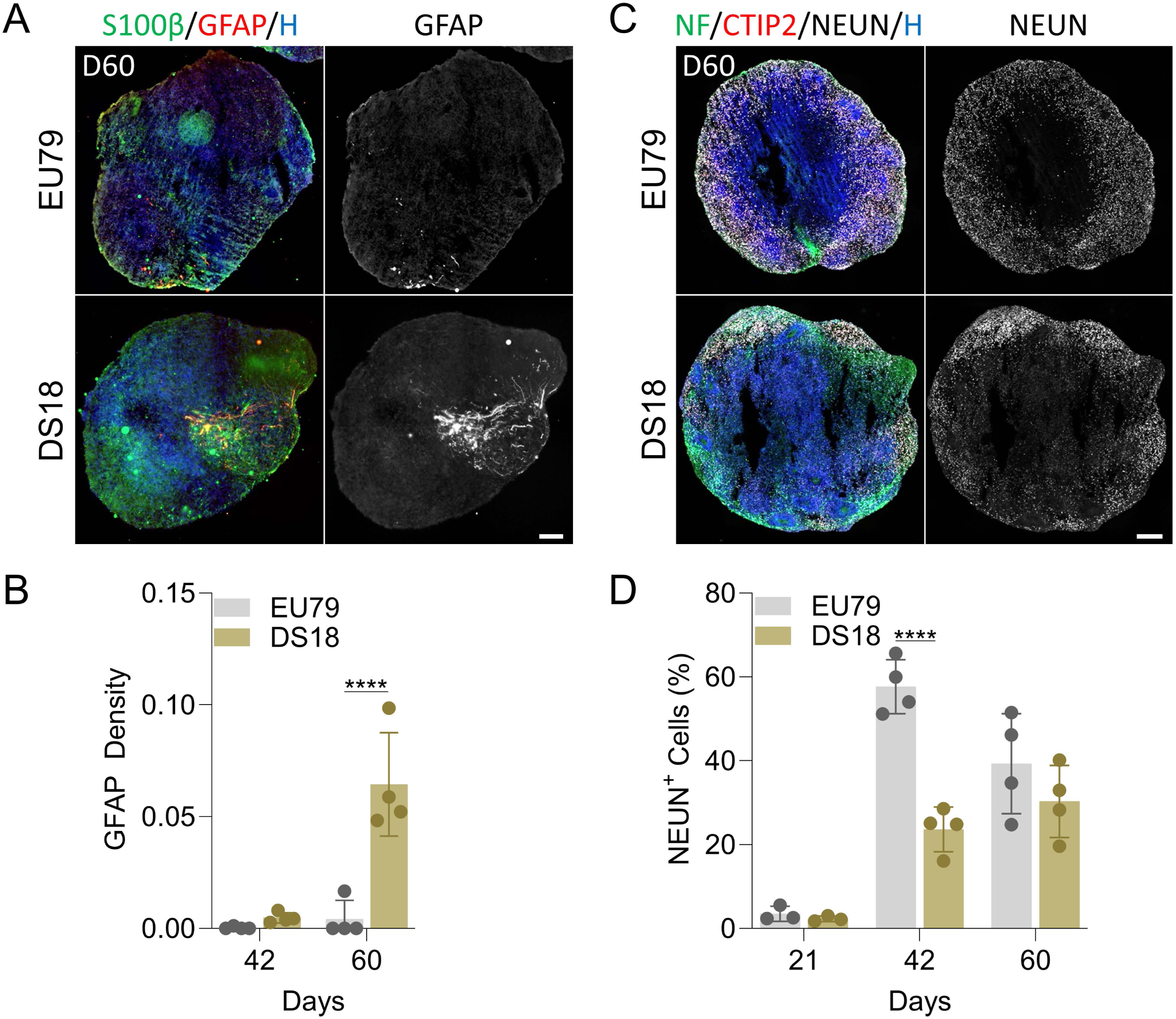
Trisomy 21 alters astroglial maturation and neuronal development in OL brain organoids. **(A)** Images of sectioned OL-brain organoids stained with S100β (green), GFAP (red) and nuclei counterstained with Hoechst (Blue) in EU79 and DS18 groups at day 60. Scale bar = 100 µm. **(B)** Bar graphs show the quantification of density of GFAP signal intensity normalised to total area at days 42 and 60 in OL-brain organoids. Data are means ± S.D.; n = 16. \*\*\*\**P* < 0.0001 via unpaired two tailed Student’s *t* test. **(C)** Images of sectioned OL-brain organoids stained with neurofilament (NF) (green), CTIP2 (Red), NEUN (Gray) and nuclei counterstained with Hoechst (Blue) in EU79 and DS18 groups at day 60. Scale bar = 200 µm. **(D)** Bar graphs show the quantification of percentage of NEUN positive cells normalised to total cells at days 21, 42 and 60 in OL-brain organoids. Data are means ± S.D.; n = 24. \*\*\*\**P* < 0.0001 via unpaired two tailed Student’s *t* test.

Taken together, these findings show that trisomic OLBOs reproduce several central features of DS-associated brain pathology, including early growth impairment, reduced oligodendroglial specification, defective oligodendrocyte maturation and myelination, increased astroglial differentiation, and delayed neuronal development. The ability of OLBOs to capture these coordinated cellular abnormalities supports their utility as a human model for dissecting the mechanisms underlying DS white matter and neurodevelopmental pathology.

### Spontaneous Hyperactivity and Impaired Evoked Network Responses in Trisomic OL-Brain Organoids

Altered excitation-inhibition balance is a recognised feature of the DS brain, although the emergence of these functional abnormalities during early human neurodevelopment remains poorly understood. To determine whether OLBOs generate functional neuronal networks suitable for electrophysiological interrogation, we first examined synaptic and neuronal subtype markers. Immunostaining of day 60 OLBO sections showed co-localisation of the presynaptic marker synaptophysin with the postsynaptic marker PSD95, consistent with synapse formation within the organoids (Fig. S3A). In parallel, both GABA-positive inhibitory neurons and GLUR3-positive excitatory neuronal populations were detected at day 60 (Fig. S3A), indicating that OLBOs contain mixed neural networks composed of relevant excitatory and inhibitory synapses. We next assessed whether these networks exhibit altered electrophysiological behaviour in trisomic organoids. OLBOs derived from the trisomic DS18 line and the euploid EU79 control line were harvested at day 60, seeded onto multi-electrode array (MEA) BioChips, and recorded after one week of attachment (Figs. 4A; process summarized in Fig. S3B). Under baseline conditions, trisomic DS18 organoids displayed significantly increased spontaneous activity compared with euploid EU79 controls, as evidenced by a higher mean firing rate and increased burst frequency, whereas burst duration was not significantly altered (Fig. 4B). These findings indicate that embryonic cortical neurons in DS brain develop a hyperactive spontaneous network state during development.

**Figure 4.**
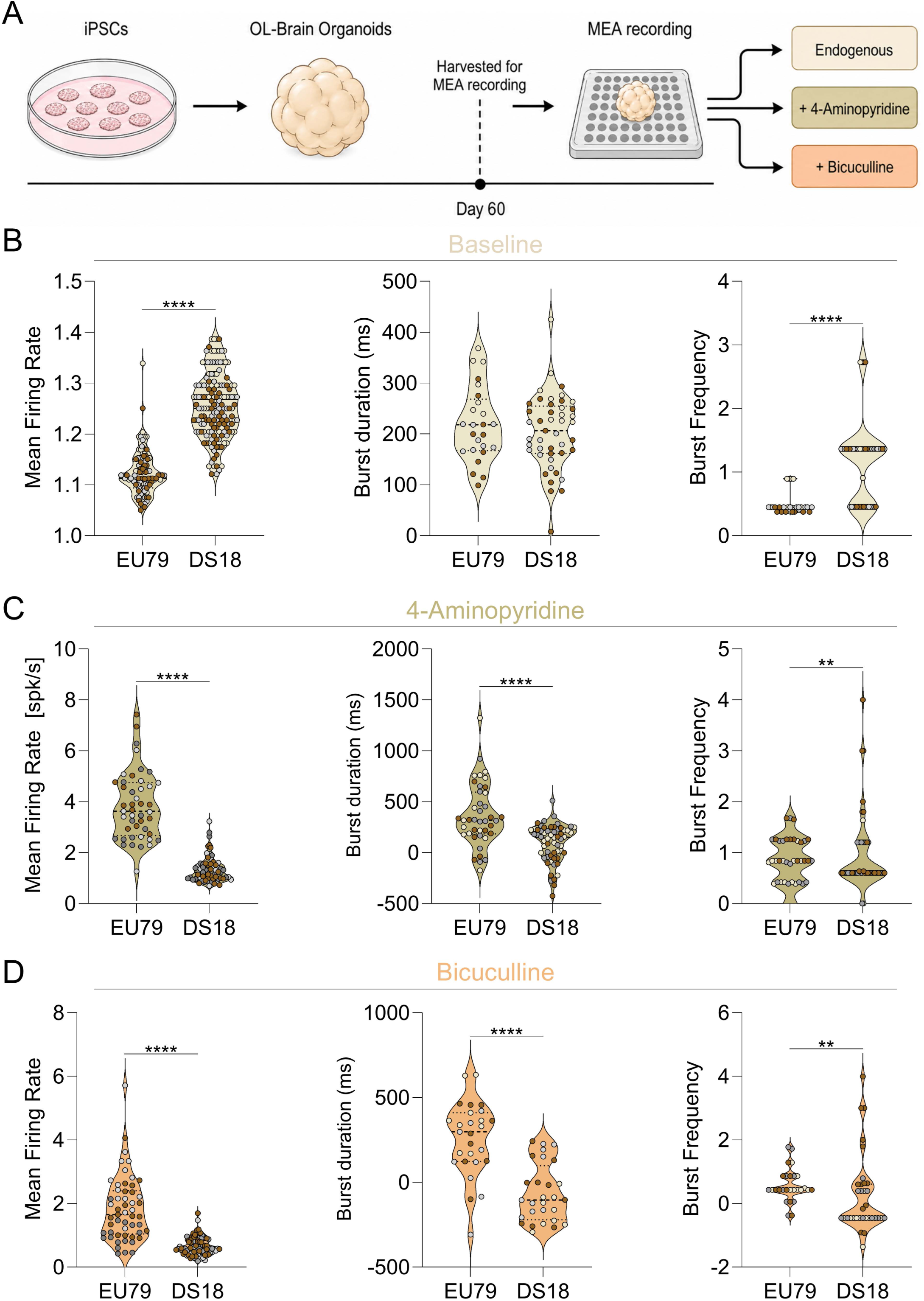
Trisomic OL brain organoids exhibit spontaneous hyperactivity but impaired pharmacological network responses. **(A)** Schematic diagram of the MEA experimental workflow. **(B)** Violin plots of baseline activity of OL-brain organoids measured parameters including mean firing rate [spk/s], burst duration (ms), and burst frequency. N = 6. \*\*\*\**P* < 0.0001 via unpaired two tailed Student’s *t* test. **(C)** Violin plots of response to 4-aminopyridine (4-AP, 100 µM) of OL-brain organoids measured parameters including mean firing rate [spk/s], burst duration (ms), and burst frequency. N = 6. \*\**P* < 0.01, \*\*\*\**P* < 0.0001 via unpaired two tailed Student’s *t* test. **(D)** Violin plots of response to bicuculline (10 µM) of OL-brain organoids measured parameters including mean firing rate [spk/s], burst duration (ms), and burst frequency. N = 6. \*\**P* < 0.01, \*\*\*\**P* < 0.0001 via unpaired two tailed Student’s *t* test.

To probe network excitability further, organoids were challenged with 4-aminopyridine (4-AP), a voltage-gated potassium channel blocker that enhances neuronal excitability and neurotransmitter release [21,22]. In euploid EU79 organoids, 4-AP elicited the expected increase in network output (Fig. 4C). By contrast, trisomic DS18 organoids exhibited a markedly blunted response, with significantly lower mean firing rate and shorter burst duration when compared to EU79 organoids (Fig. 4C). Notably, burst frequency remained higher in DS18 organoids despite the reduction in overall firing output and burst length (Fig. 4C), suggesting that trisomic networks are unable to sustain coordinated bursting and instead display fragmented or inefficient activity patterns following stimulation. We next challenged the organoids with bicuculline, a GABA_A_ receptor antagonist that increases network activity through disinhibition [23]. As observed with 4-AP, bicuculline-treated DS18 organoids showed significantly reduced mean firing rate and shorter burst duration relative to euploid EU79 controls, while burst frequency was increased (Fig. 4D). Thus, although trisomic DS18 organoids display enhanced spontaneous bursting, they fail to present a normal coordinated response to pharmacological stimulation.

### Transcriptomic Profiling Reveals Early Developmental Dysregulation and Late Synaptic Imbalance in Trisomic OL-brain Organoids

To begin to define the molecular programs underlying the developmental and functional abnormalities observed in trisomic organoids, we performed bulk RNA sequencing on DS18 and EU79 OL-brain organoids collected at days 35, 42, and 60 of differentiation (Fig. 5A). Unsupervised clustering revealed clear transcriptomic divergence between trisomic DS18 and euploid EU79 organoids across development (Fig. S4A). Principal component (PC) analysis further separated samples according to both genotype and developmental stage, with PC1 and PC2 explaining 40.33% and 22.92% of the total variance, respectively (Fig. 5B). Day 35 and day 42 trisomic DS18 samples clustered far from their euploid EU79 counterparts indicating genotype difference, while day 60 samples occupied a distinct position indicating less genotype difference likely driven by maturation-related transcriptional shift in both groups as brain develops.

**Figure 5.**
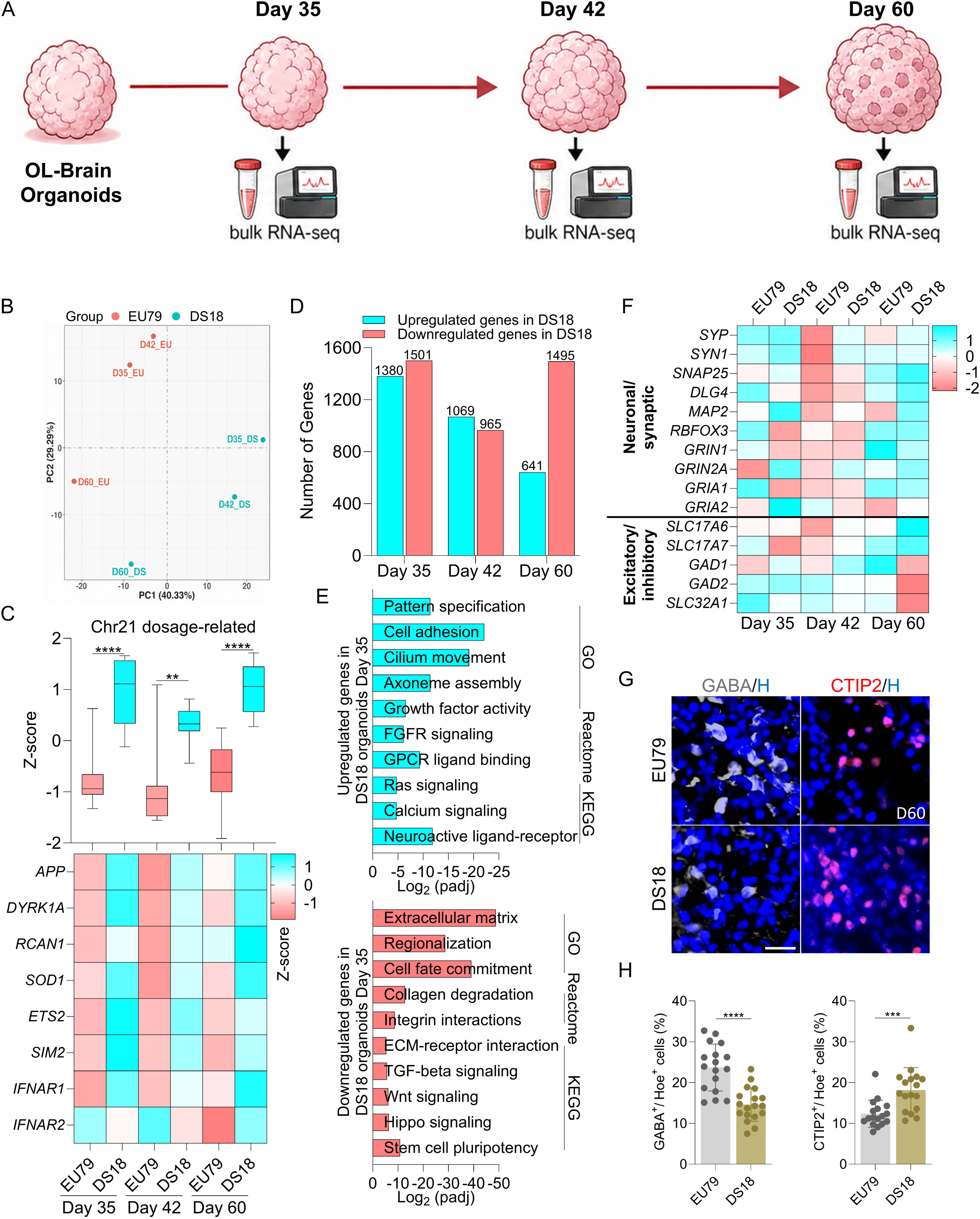
Transcriptomic profiling reveals chromosome 21 dosage effects, disrupted developmental pathways, and excitatory inhibitory imbalance in trisomic OL brain organoids. **(A)** Schematic diagram of bulk RNA-sequencing experimental workflow showing timeline of EU79 and DS18 OL-brain organoids harvested at days 35, 42, and 60. **(B)** Principal component analysis of RNA-seq read counts derived from euploid EU79 and trisomic DS18 OL-brain organoids at days 35, 42 and 60. **(C)** Box blot showing distribution of Chromosome 21 genes (listed in heatmap) obtained from bulk RNA-seq of EU79 and DS18 OL-brain organoids at days 35, 42, and 60. Data minimum to maximum; \*\**P* < 0.01,\*\*\*\**P* < 0.0001 via one-way ANOVA. Below heatmap of Chromosome 21 genes, values are shown as *z* score. **(D)** Bar graph showing the number of significantly up- (cyan) and down-regulated (red) differentially expressed genes (DEGs) in DS18 versus EU79 at days 35, 42, and 60. **(E)** Bar graphs showing gene ontology (GO), reactome and KEGG enrichment analysis of upregulated genes (cyan), downregulated genes (red), in 35 days OL-brain organoids derived from DS18 compared with EU79 organoids. **(F)** Heatmap (Z-score) of genes associated with neuronal/synaptic and excitatory/inhibitory neurons across development of OL-brain organoids derived from DS18 and EU79 groups **(G)** Images of sectioned OL-brain organoids stained with GABA (Gray), CTIP2 (Red), and nuclei counterstained with Hoechst (Blue) in EU79 and DS18 groups at day 60. Scale bar = 20 µm. **(H)** Bar graphs show the quantification of percentage of GABA positive cells and CTIP2 positive cells normalised to total Hoechst positive cells at day 60 in OL-brain organoids. Data are means ± S.D.; n = 15. \*\*\**P* < 0.001,\*\*\*\**P* < 0.0001 via unpaired two tailed Student’s *t* test.

To confirm the expected dosage effect of trisomy 21, we next examined the expression of chromosome 21-associated genes. Z score analysis demonstrated consistent enrichment of canonical HSA21 dosage-sensitive transcripts, including A*PP, DYRK1A, RCAN1, SOD1, ETS2, SIM2, IFNAR1*, and *IFNAR2*, in DS18 organoids at all three time points relative to EU79 controls (Fig. 5C). This confirmed preservation of the trisomic transcriptional signature throughout organoid development. Differential expression analysis (*see methods*) revealed marked stage-dependent dysregulation in DS18 organoids, with the largest burden of differentially expressed genes was detected at day 35, with 1380 upregulated and 1501 downregulated genes in DS18 relative to EU79 (Fig. 5D; Fig. S4B). At day 42, the number of dysregulated genes decreased to 1069 upregulated and 965 downregulated, whereas at day 60 the transcriptional profile shifted toward predominant repression, with 641 upregulated and 1495 downregulated genes (Figs. 5D, S4B). These findings indicate that the major transcriptomic perturbation emerges early during lineage specification and persists into later maturation stages.

To determine whether trisomy 21 dysregulated a stable or temporally evolving set of genes, we compared the overlap among genes upregulated in DS18 across the three developmental stages. Most upregulated genes were stage-specific, with 865 genes unique to day 35, 543 unique to day 42, and 398 unique to day 60 (Fig. S4C). Only 111 genes were shared across all three stages, whereas 343, 60, and 72 genes were shared between day 35/day 42, day 35/day 60, and day 42/day 60, respectively (Fig. S4C). Thus, trisomy 21 does not impose a static transcriptional state, but rather drives a temporally dynamic dysregulation of developmental programs.

Because the strongest molecular divergence was observed at day 35, we next performed pathway enrichment analysis on day 35 differentially expressed genes. Genes upregulated in DS18 organoids were enriched for processes related to pattern specification, cell adhesion, cilium movement, axoneme assembly, growth factor activity, FGFR signaling, Ras signaling, calcium signaling, GPCR ligand binding, and neuroactive ligand-receptor interaction (Fig. 5E, top panel). By contrast, genes downregulated in DS18 were enriched for pathways associated with extracellular matrix organisation, regionalisation, cell fate commitment, collagen degradation, integrin interactions, ECM-receptor interaction, TGF-beta signaling, Wnt signaling, Hippo signaling, and stem cell pluripotency (Fig. 5E, lower panel). These findings suggest that trisomic OL-brain organoids undergo early disruption of morphogenetic, cell fate, and matrix-associated programs, which may precede the structural and functional abnormalities detected at later stages.

To relate the day 60 transcriptomic changes to the functional phenotype, we next examined genes associated with neuronal identity, synaptic function, and excitatory/inhibitory specification. This analysis indicated relative enrichment of neuronal and synaptic transcripts in DS18 organoids at day 60, together with increased expression of excitatory neuronal markers and reduced expression of inhibitory markers (Fig. 5F). We then validated these findings at the protein level by immunostaining for the inhibitory neuronal marker GABA and the excitatory cortical neuronal marker CTIP2. Consistent with the transcriptomic data, DS18 organoids showed a significant reduction in GABA-positive cells and a significant increase in CTIP2-positive cells compared with EU79 controls (Fig. 5G). These findings support a shift in neuronal composition toward excitatory identity in trisomic DS18 organoids and are consistent with the spontaneous network hyperactivity and abnormal pharmacological responses detected by MEA at the same stage (Fig. 4).

Collectively, these data show that trisomic OL-brain organoids undergo pronounced and dynamic transcriptional dysregulation across development, with the strongest divergence emerging during early lineage specification. Early disruption of patterning, fate commitment, and extracellular matrix programs is followed by later alterations in neuronal and synaptic gene expression, providing a molecular framework for the cellular and electrophysiological abnormalities observed in the trisomic organoids.

## Discussion

In this study, we used OL-brain organoids to investigate early cellular, functional, and molecular abnormalities associated with Down syndrome during human neurodevelopment. Although trisomy 21 is the established genetic cause of Down syndrome, the developmental sequence linking this chromosomal imbalance to later white matter defects, glial abnormalities, and neuronal dysfunction remain incompletely defined. Here, trisomic organoids reproduced several key features of Down syndrome neuropathology, including early growth delay, impaired neural cells output, spontaneous network hyperactivity with abnormal pharmacological responses, and marked stage dependent transcriptomic dysregulation. Together, these findings support the view that major pathological features of the Down syndrome brain arise early during neural and glial specification.

A major strength of this study was the use of a human 3D organoid system enriched for oligodendroglial lineage cells while preserving parallel development of neurons and astrocytes. Previous studies of Down syndrome brain pathology have relied on postmortem tissue [2,24], animal models [6], or 2D iPSC derived cultures [8], each of which has clear limitations in modelling early human development. By contrast, the OL-brain organoid platform used here enables interrogation of multiple neural lineages within a shared developmental context. The use of an isogenic disomic control further strengthens the model by minimising the effects of genetic background. Consistent with our previous work [12], this system reproducibly generated oligodendroglial cells together with maturing neurons and astrocytes, providing a suitable framework to examine the developmental consequences of trisomy 21. The first major finding was that trisomic organoids showed impaired early growth, with reduced size during the initial stages of differentiation, consistent with previous studies [9], followed by partial catch up at later stages. These findings suggest that trisomy 21 exerted its strongest effects during early proliferation, patterning, or lineage progression rather than on gross tissue expansion alone. This interpretation was supported by the cellular data, where trisomic DS18 organoids showed reduced early DCX expression and delayed accumulation of NEUN positive neurons, indicating impaired neuronal differentiation. Thus, even where overall organoid size later approached that of controls, lineage specific defects persisted, suggesting that apparent morphological recovery does not reflect normal developmental progression.

A central outcome of this study was the demonstration of defective oligodendroglial maturation in trisomic organoids. Although SOX10 positive cells emerged in both groups, trisomic organoids exhibited reduced SOX10 expression during early differentiation and markedly reduced PLP1 and MBP expression at later stages. These findings indicate that oligodendroglial specification was initiated in trisomic DS18 organoids but efficient progression toward mature myelinating oligodendrocytes was compromised. These observations are in agreement with previous studies in Down syndrome mouse models and human mid-fetal brain showing defects in oligodendrocyte differentiation and myelin production [6]. Our data extend those observations by showing that such abnormalities are detectable in a human organoid system during early developmental stages. The partial recovery in SOX10 positive cell numbers by day 60, despite persistent deficits in maturation markers, further suggest that delayed timing alone cannot fully explain the phenotype and that maturation itself was impaired. In parallel, trisomic organoids showed a significant increase in GFAP positive astrocytes at later stages. This finding was consistent with previous reports describing increased astrogliogenesis in Down syndrome [2,20]. In the context of the present model, the combination of reduced mature oligodendrocyte markers and increased astrocytes suggests altered glial lineage balance. Given the developmental plasticity of glial progenitors [25], one possibility is that OPCs are diverted toward an astroglial fate under trisomic conditions. While this cannot be concluded directly from the current data, the cellular and transcriptomic findings together support a shift in glial output that favours astrocytic differentiation over oligodendroglial maturation. Future lineage tracing approaches will be required to resolve this directly.

At the functional level, trisomic organoids displayed spontaneous network hyperactivity but impaired evoked responses. OL-brain organoids contained synaptic structures together with excitatory and inhibitory neuronal populations, supporting the formation of functional neural networks. MEA recordings showed increased baseline firing and burst frequency in trisomic organoids, indicating a hyperactive spontaneous state. However, following treatment with 4-aminopyridine or bicuculline, trisomic organoids failed to mount a coordinated response comparable to controls, showing reduced firing output and shorter burst duration despite persistent high burst frequency. These findings suggest that trisomic networks are not simply hyperexcitable, but functionally disorganised and less able to sustain coordinated activity. Such a phenotype is consistent with immature or unstable circuit formation, in line with previous studies showing that trisomy 21 disrupts excitatory/inhibitory balance and neuronal network communication in cerebral organoids, and that early DS neuronal networks exhibit impaired synchrony and altered synaptic excitability [26,27]. Interestingly, the direction of excitatory and inhibitory imbalance observed here differs from some previous studies in more mature Down syndrome systems [28], where increased inhibitory drive has often been emphasised [29]. In our organoids, transcriptomic and protein level analyses at day 60 supported relative enrichment of excitatory neuronal markers together with depletion of inhibitory markers. This may also relate to the developmental stage and regional identity captured by the organoids. During normal early brain development, neural circuits are initially biased toward excitation, whereas inhibitory control is progressively established as circuits mature and behavioural outputs become more refined [30]. Because our organoids model an early cortical developmental window and are not designed to robustly generate ventral forebrain interneuron populations that normally migrate into the cortex, the relative enrichment of excitatory over inhibitory neuronal features observed here may reflect an immature network state rather than the final maturation of the DS brain.

This discrepancy may reflect the early developmental stage captured by the present model. It may also link to the regional identity of the organoids, which are not designed to robustly generate ventral forebrain interneuron populations that normally migrate into the cortex. Thus, the imbalance observed here may represent an early developmental state rather than the final maturation of the Down syndrome brain. Nevertheless, these data suggest that altered network assembly begins early and may precede later circuit level abnormalities.

The transcriptomic analysis provided a molecular framework for these developmental and functional phenotypes. Trisomic and euploid organoids separated clearly across all stages, with the highest burden of differentially expressed genes at day 35. This is notable because it places the major molecular divergence before the overt electrophysiological phenotype observed at day 60. Pathway enrichment at day 35 indicated dysregulation of receptor signalling, ciliary and axonemal programs, and neuroactive ligand receptor pathways among upregulated genes, whereas downregulated genes were enriched for extracellular matrix organisation, cell fate commitment, and Wnt, Hippo, and TGF beta signalling. These findings point to early disruption of tissue organisation and developmental signalling programs that are likely to influence both glial and neuronal maturation. By day 60, transcriptomic changes were more strongly linked to neuronal and synaptic function, including altered excitatory and inhibitory signatures, consistent with the immunostaining and MEA data. Together, these findings support the concept that white matter and circuit dysfunction in Down syndrome have developmental origins and begin during early neural and glial specification.

This study has limitations. The analysis was performed using one trisomic line and one isogenic euploid control, and validation in additional lines will be important. In addition, the organoid system does not fully recapitulate all anatomical domains, or extrinsic inputs present in the developing human brain. Finally, although our data strongly support altered glial lineage balance and delayed neural maturation, longer term culture and lineage tracing studies will be required to distinguish delayed maturation from persistent arrest and to define the origin of the astroglial increase more precisely.

## Materials and Methods

### Human iPSC culture

Engineered mMaple (SOX10-GFP) WTC iPSC line [13], together with the DS18 and EU79 iPSC lines [8], were maintained on Matrigel coated plates (StemCell Technologies, Cat. #354277) in mTeSR Plus medium (Stem Cell Technologies, Cat. #100-0276) according to the manufacturer’s instructions. Cells were cultured under standard humidified conditions at 37 °C with 5% CO₂ and were routinely passaged to maintain pluripotency and culture quality.

### Generation of Oligodendrocyte Enriched Brain Organoids

Oligodendrocyte enriched brain organoids were generated as previously described [13,23] with minor modifications. Briefly, human iPSCs were differentiated into 2D human neuroectodermal colonies by dual SMAD inhibition using 10 µM SB431542 (Sapphire Biosciences, Cat. #A10826) and 0.1 µM LDN193189 dihydrochloride (Sigma, Cat. #SML0559) for 3 days in N2 medium. N2 medium consisted of DMEM/F12 (Gibco, Cat. #11320-33), 2% B27 supplement (Gibco, Cat. #17504044), 1% N2 supplement (Gibco, Cat. #17502-048), 1% MEM non-essential amino acids (Gibco, Cat. #11140-050), 1% penicillin/streptomycin (Gibco, Cat. #15140148), and 0.1% β mercaptoethanol (Gibco, Cat. #21985-023) [31]. Human neuroectodermal colonies were detached using dispase at 37 °C for 20 min and transferred to 6 well ultra-low attachment plates to form neuroepithelial spheroids. Spheroids were maintained for 4 days in Olig2 medium supplemented daily with 40 ng/mL bFGF to promote neural stem cell proliferation and self-organization. Olig2 medium consisted of DMEM/F12 (Thermo Fisher Scientific, Cat. #11320033), 2% B27 supplement without vitamin A (Invitrogen, Cat. #12587-010), 1% N2 supplement (Thermo Fisher Scientific, Cat. #17502001), 1% MEM non-essential amino acids (Thermo Fisher Scientific, Cat. #11140050), 1% penicillin/streptomycin (Invitrogen, Cat. #15140122), 0.1% 2-mercaptoethanol (Thermo Fisher Scientific, Cat. #21985023), 10 ng/mL human IGF-I (Capsugel, Cat. #100-11-1000), 10 ng/mL human NT-3 (Capsugel, Cat. #450-03-100UG), 60 ng/mL 3,3′,5-triiodo-L-thyronine (T3; Sigma-Aldrich, Cat. #T2877-250MG), 10 ng/mL human HGF (Capsugel, Cat. #100-39H-100), 100 ng/mL biotin (Sigma-Aldrich, Cat. #B4639-1G), 10 µM cAMP (Sigma-Aldrich, Cat. #D0627), and 10 ng/mL PDGF-AA (Capsugel, Cat. #100-13A-100UG). To generate organoids, neuroepithelial spheroids were transferred to 96 well ultra-low attachment plates and differentiated using the free embedding method in Olig3 medium containing 1% Matrigel (StemCell Technologies, Cat. #354277) [12]. The Olig3 medium consisted of DMEM/F12 (Thermo Fisher Scientific, Cat. #11320033) and Neurobasal medium (Invitrogen, Cat. #21103049) in a 1:1 ratio, 1% B27 supplement without vitamin A (Invitrogen, Cat. #12587-010), 0.5% N2 supplement (Thermo Fisher Scientific, Cat. #17502001), 1% GlutaMAX (Thermo Fisher Scientific, Cat. #35050061), 1% MEM non-essential amino acids (Thermo Fisher Scientific, Cat. #11140050), 1% penicillin/streptomycin (Invitrogen, Cat. #15140122), 0.1% 2-mercaptoethanol (Thermo Fisher Scientific, Cat. #21985023), 10 ng/mL human IGF-I (Capsugel, Cat. #100-11-1000), 2.5 µg/mL insulin (Sigma-Aldrich, Cat. #I9278-5ML), 10 ng/mL human NT-3 (Capsugel, Cat. #450-03-100UG), 60 ng/mL 3,3′,5-triiodo-L-thyronine (T3; Sigma-Aldrich, Cat. #T2877-250MG), 10 ng/mL human HGF (Capsugel, Cat. #100-39H-100), 100 ng/mL biotin (Sigma-Aldrich, Cat. #B4639-1G), 10 µM cAMP (Sigma-Aldrich, Cat. #D0627), and 10 ng/mL PDGF-AA (Capsugel, Cat. #100-13A-100UG). After 3 days, organoids were transferred to 24 well ultra-low attachment plates. Medium was replaced three times per week throughout differentiation.

### Immunostaining

***Whole mount immunostaining*.** Whole mount immunostaining was performed as previously described [32]. Organoids were fixed in 4% paraformaldehyde in PBS for 20 min at room temperature, washed three times for 10 min in PBS, and permeabilized in 0.1% Triton X 100 in PBS (PBST). Samples were blocked overnight at room temperature in 6% bovine serum albumin (BSA; Sigma, Cat. #A9418-50G) in PBST. Organoids were then incubated with primary antibodies (Table 1) for 48 h at 4 °C, washed three times in PBST, and incubated with the appropriate Alexa Fluor conjugated secondary antibodies overnight at room temperature. Both primary and secondary antibodies were diluted in PBST (see Table 1 for dilutions). After secondary incubation, samples were washed three times in PBST and cleared in a solution containing 25% urea and 65% sucrose in water. All steps were done using an orbital shaker. Organoids were mounted in double concave slides and counterstained with Hoechst 33342. Images were acquired using a Leica SP8 point scanning confocal microscope. The staining and imaging settings were the same across the two genotypes.

**Table 1:** List of Antibodies used for immunostaining.

| Antigen | RRID Number | Source | Cat# | Dilution |  |
| --- | --- | --- | --- | --- | --- |
|  |  |  |  | IHC | WM |
| CNPase | AB_476854 | Sigma | C5922 | 1:300 | - |
| DCX | AB_10610966 | Santa Cruz | sc-271390 | 1:500 | 1:150 |
| GABA | N.A. | ThermoFisher | 483900 | 1:300 | - |
| GFAP | N.A. | ThermoFisher | 71-1500 | 1:500 | 1:150 |
| GFP | AB_300798 | Abcam | ab13970 | 1:500 | 1:150 |
| GLUR3 | AB_86916 | ThermoFisher | 32-0400 | 1:300 | - |
| MBP | AB_2799920 | Cell signaling | 78896 | 1:300 | - |
| NCAD | AB_2798427 | Cell Signaling | 14215 | 1:300 | 1:150 |
| NEUN | AB_2298772 | Milipore | MAB377 | 1:500 | 1:150 |
| NF | AB_477272 | Sigma-Aldrich | N4142 | 1:1000 | 1:150 |
| PLP1 | AB_10737663 | Sigma | SAB1404221 | 1:300 | - |
| S100β | AB_882426 | Abcam | ab52642 | 1:300 | 1:150 |
| SOX10 | N.A. | Cell Signaling | 89356 | 1:300 | 1:150 |
| CTIP2 | AB_2064130 | Abcam | ab18465 | 1:500 | - |

***Immunohistochemistry*.** Immunohistochemistry was performed as previously described [33]. Organoids were fixed overnight in 4% paraformaldehyde in PBS at room temperature, washed three times in PBS, and cryoprotected in 30% sucrose in PBS at 4 °C until fully equilibrated. Samples were then embedded in a 3:2 mixture of optimal cutting temperature compound and 30% sucrose on dry ice. Cryosections of 16 µm thickness were collected onto Superfrost™ Plus Microscope Slides (Thermo Scientific, Cat. #SF41296). For staining, sections were washed three times in PBS for 10 min each, blocked for 1 to 6 h at room temperature in 3% BSA and 0.1% Triton X 100 in PBS (PBST), and incubated overnight at 4 °C with primary antibodies (Table 1). After three washes in PBS, sections were incubated with the appropriate Alexa Fluor conjugated secondary antibodies for 1 h at room temperature, counterstained with Hoechst 33342, and mounted for imaging. Antibody dilutions were in blocking solution as per Table 1. Images were acquired using a Zeiss Axio Scan Z1 fluorescent imager. The staining and imaging settings were the same across the two genotypes. Alexa Fluor 488, 546, and 633 secondary antibodies were obtained from Thermo Fisher Invitrogen (Table 1).

### Western blotting

Organoids were collected and lysed by sonication in RIPA buffer (Thermo Scientific, Cat. #89900) supplemented with protease inhibitor cocktail (20 µL per 1 mL lysis buffer). Protein concentration was determined using the Pierce BCA Protein Assay Kit (Pierce Biotechnology, Rockford, IL, USA). Equal amounts of total protein (20 to 30 µg) were loaded and separated by 10% SDS PAGE and transferred onto nitrocellulose membranes. Membranes were blocked in 5% skim milk in 1× TBST (Tris-Buffered Saline with Tween 20) for 1 h at room temperature and then incubated overnight at 4 °C with primary antibodies diluted 1:1000 in 3% bovine serum albumin in 1× TBST. After washing three times in 1× TBST, membranes were incubated with the appropriate secondary antibodies diluted 1:10000 in 5% skim milk in 1× TBST for 1 h at room temperature. Protein bands were visualised using enhanced chemiluminescence reagents (Thermo Scientific, Pittsburgh, PA, USA). Membranes were subsequently stripped and re-probed with anti-actin antibody as a loading control. Band intensity was quantified using ImageJ [34].

### Microelectrode Array Recordings

MEA experiments were performed using the 3Brain dissociated cultures protocol with modifications (Fig. S3B). Briefly, BioChips were rehydrated with sterile distilled water for 1 to 2 h, sterilized by external cleaning with absolute ethanol and internal incubation with 70% ethanol for 20 min, rinsed 4 to 5 times with sterile distilled water, and dried under a laminar flow hood. Chips were then incubated overnight with sterile medium to enhance hydrophilicity and improve cell attachment. For surface coating, BioChips were incubated with 0.1 mg/mL poly L ornithine, rinsed thoroughly with sterile distilled water, and then coated with 50 µg/mL laminin at 37 °C for 4 h. After removal of the coating solution, organoids were placed directly onto the coated electrode surface in a small drop of medium. After 1 h of attachment, the reservoir was filled with approximately 2.5 mL of Olig3 medium and maintained at 37 °C in 5% CO₂ and 95% air. Recordings were performed after at least 1 week of attachment. One third of the medium was replaced weekly to compensate for evaporation and maintain culture stability. Electrical activity was recorded using BrainWave4 software and analysed using BrainWave5 software (3Brain). For pharmacological stimulation, 4 aminopyridine was prepared in 100 µL medium at a final concentration of 100 µM and incubated for 10 min at 37 °C before recording. Bicuculline was prepared in 100 µL medium at a final concentration of 10 µM and added directly during recording. Because bicuculline stock was prepared in dimethyl sulfoxide, vehicle control recordings were performed by adding an equivalent volume of DMSO to exclude solvent related effects on neuronal activity.

### RNA Extraction and Bulk RNA Sequencing

Whole organoids were snap frozen and stored at −80 °C until RNA extraction. Total RNA was isolated using the NucleoSpin RNA Mini kit (Macherey Nagel, Cat. #740955.50) according to the manufacturer’s instructions, as previously described [35]. RNA concentration and purity were assessed using a NanoDrop 1000 spectrophotometer (Thermo Scientific). Samples were submitted to Novogene Co., Ltd. for library preparation, sequencing, and primary bioinformatic processing [36].

***Library preparation, sequence alignment, and differential expression analysis.*** Messenger RNA was enriched from total RNA using poly T oligo attached magnetic beads. Following fragmentation, first strand cDNA was synthesized using random hexamer primers, followed by second strand synthesis. Libraries were prepared using standard end repair, A tailing, adaptor ligation, size selection, amplification, and purification steps. Library quality was assessed using Qubit, real time PCR, and Bioanalyzer analysis before sequencing on an Illumina platform to generate paired end reads. Raw FASTQ files were processed using fastp to remove adaptor sequences, reads containing poly N, and low-quality reads. Clean reads were aligned to the reference genome using HISAT2 v2.0.5. Gene level read counts were generated using featureCounts v1.5.0-p3, and expression levels were calculated as fragments per kilobase of transcript per million mapped reads. Differential gene expression analysis was performed using edgeR v3.22.5 following normalization with a scaling factor for each library. P values were adjusted using the Benjamini Hochberg method [37]. Genes with an adjusted P value < 0.05 and an absolute fold change ≥ 2 were considered significantly differentially expressed.

***Functional enrichment analysis.*** Gene Ontology and pathway enrichment analyses of differentially expressed genes were performed using the ShinyGO application [38]. Enrichment analysis was conducted separately for each developmental stage and for upregulated and downregulated gene sets as indicated in the Results.

### Statistical analysis

Data are presented as mean ± standard deviation (S.D.) unless otherwise indicated. For non-normally distributed data determined by employing a Shapiro-Wilk test for normality, median values were used as specified in the relevant figure legends. The number of biological replicates (*n*) and sample size for each experiment are provided in the figure legends, with a minimum of three biological replicates used in this study. ImageJ was used for quantification of signal intensity and positive cell number from microscopy images. For comparison between two groups, unpaired two tailed Student’s t tests were used. For comparisons involving more than two groups, one-way ANOVA or two-way ANOVA was used as appropriate. Statistical analyses were performed using GraphPad Prism v10, and *P* < 0.05 was considered statistically significant.

## Acknowledgments

B.A.-M. and S.D.M. acknowledge the University of Queensland (UQ) Research Training Scholarship. B.A.-M. also would like to thank the UQ Entrepreneurial PhD Top-up Scholarship. We gratefully acknowledge Bruce Conklin (Department of Medicine, Gladstone Institute of Cardiovascular Disease) for generously providing the WTC iPSC lines, and the School of Biomedical Sciences’ Microscopy and Image Analysis Facility UQ.

## Author Contributions

M.B.M. performed and designed experiments, analysed data, interpreted results, and drafted the manuscript. S.D.M., B.A.-M., S.P., performed additional experiments. P.G.N., contributed to supervision and experimental discussions. E.J.W. and M.R.S. conceived and supervised the study, interpreted results, and co-wrote the manuscript. Critical feedback was shared and discussed by all authors, leading to all authors reviewed and approving the final version of the manuscript.

## Funding

M.R.S. is supported by the Start-up Funding (SF-2025_003), Interdisciplinary Research Program (IDRP-2019-001), and Qatar Research, Development, and Innovation Council (ANMR01-0209-250025). E.J.W. is supported by the Australian National Health and Medical Research Council (NHMRC) through grants NHMRC 2020434, MRFF 2024380 and GA56111. P.G.N. is supported by School of Biomedical Sciences UQ and by grant FIGHT MND DIS-202403-01216.

## Data Availability Statement

The data that support the findings of this study are available from the corresponding author upon reasonable request.

## Declarations

### Ethics approval and consent to participate

All experiments were conducted in compliance with the ethical guidelines of the University of Queensland and approved by the University of Queensland Human Research Ethics Committee (Approval number 2019000159).

## Consent for publication

Not applicable.

## Conflict of Interest

The authors declare no competing interests.

## List of Tables Table 1: List of Antibodies used for immunostaining

* WM: Wholemount; N.A.: Not Available

## Supplementary Figure Legend

**Figure S1. Extended validation of growth dynamics and lineage specification in OL brain organoids.**

**(A)** Images showing the developmental stages of OL-brain organoids derived from DS18 and EU79 iPSCs over time in culture *in vitro* under brightfield, and green fluorescence protein (GFP). GFP indicates SOX10 positive cells. Right panel is graph showing the growth difference between euploid and trisomic DS18 OL-brain organoids (based on the average diameter) at different stages of *in vitro* culture. Data are presented as the mean ± standard deviation (*n* = 3). The total number of analyzed organoids is 80. Scale bars = 200-900 µm;
**(B)** Images of sectioned OL-brain organoids stained with SOX10 (Gray), GFP (Green), and nuclei counterstained with Hoechst (Blue) in mMaple WTC iPSC line at days 7, 35, 42, and. Scale bars = 100 µm; magnified, 20 µm.
**(C)** Images of sectioned OL-brain organoids stained with CNPase (Yellow), MBP (Red), and nuclei counterstained with Hoechst (Blue) in mMaple WTC iPSC at day 60. Scale bar = 20 µm.
**(D)** Images of sectioned OL-brain organoids stained with S100β (Gray) and GFP (Green) in mMaple WTC iPSC line at days 7, 35, and 60, and. Scale bars = 100 µm; magnified, 20 µm.
**(E)** Immature neuronal DCX positive cells labelled in sectioned OL-brain organoids and counterstained the nuclei with Hoechst (Blue) in mMaple WTC iPSC line at days 35 and 60. Scale bars = 100 µm; magnified, 20 µm.
**(F)** Post-mitotic neuronal cells labelled with NEUN (Red) and neurofilament (NF) (Gray) in sectioned OL-brain organoids at day 21. Scale bar = 50 µm.

**Figure S2. Early neurogenic deficits and delayed neuronal maturation in trisomic OL brain organoids.**

**(A)** Images of sectioned OL-brain organoids stained with DCX (Red) and counterstained the nuclei with Hoechst (Blue) in EU79 and DS18 groups at days 10 and 60. Scale bars = 100 µm; magnified, 20 µm.
**(B)** Bar graphs show the quantification of density of DCX density normalised to total area of organoids at day 21 in OL-brain organoids. Data are means ± S.D.; n = 6. \**P* < 0.05 via unpaired two tailed Student’s *t* test.
**(C)** Staining of NEUN (Red) positive mature neurons in sections of OL-brain organoids derived from EU79 and DS18 iPSCs at days 14, 21, 35, and 42. All groups were counter stained the nuclei with Hoechst (Blue). Scale bars = 100 µm; magnified, 20 µm.

**Figure S3. Synaptic marker expression and MEA preparation workflow for OL brain organoids.**

**(A)** Staining of Synaptophysin (Red), PSD95 (Yellow), GABA (Brown) and GLUR3 (Gray) in sections of OL-brain organoids derived from mMaple WTC iPSC line at day 60. All groups were counter stained the nuclei with Hoechst (Blue). Scale bars = 20 µm.
**(B)** Overview of the ten-step workflow for MEA BioChip preparation, coating, and organoid attachment. Organoids are attached for ≥1 week before recording.

**Figure S4. Stage dependent transcriptomic dysregulation and pathway enrichment in trisomic OL brain organoids.**

**(A)** Unsupervised hierarchical clustering heatmap expression patterns of EU79 and DS18 OL-brain organoids at days 35, 42, and 60.
**(B)** Volcano plot highlighting differentially expressed genes in OL-brain organoids of DS18 vs EU79 group at days 35, 42 and 60. Red colour defines an upregulated expression with a log_2_ (fold change) > 2, green defines a downregulated expression with a log_2_ (fold change) < 2. Blue colour defines non-differentially expressed genes.
**(C)** Three-way Venn diagram of genes upregulated in DS18 across days 35, 42, and 60, with absolute counts and percentages for each stage-specific and overlapping set.
**(D)** Gene Ontology Biological Process enrichment at day 35 (top), day 42 (middle), and day 60 (bottom); x-axis, −log₁₀(FDR).

## Notes

### Competing Interest Statement

The authors have declared no competing interest.

